# Balanced bidirectional optogenetics reveals the causal impact of cortical temporal dynamics in sensory perception

**DOI:** 10.1101/2024.05.30.596706

**Authors:** Daniel Quintana, Hayley Bounds, Julia Veit, Hillel Adesnik

## Abstract

Whether the fast temporal dynamics of neural activity in brain circuits causally drive perception and cognition remains one of most longstanding unresolved questions in neuroscience^1–6^. While some theories posit a ‘timing code’ in which dynamics on the millisecond timescale is central to brain function, others instead argue that mean firing rates over more extended periods (a ‘rate code’) carry most of the relevant information. Existing tools, such as optogenetics, can be used to alter temporal structure of neural dynamics^7^, but they invariably change mean firing rates, leaving the interpretation of such experiments ambiguous. Here we developed and validated a new approach based on balanced, bidirectional optogenetics that can alter temporal structure of neural dynamics while mitigating effects on mean activity. Using this new approach, we found that selectively altering cortical temporal dynamics substantially reduced performance in a sensory perceptual task. These results demonstrate that endogenous temporal dynamics in the cortex are causally required for perception and behavior. More generally, this new bidirectional optogenetic approach should be broadly useful for disentangling the causal impact of different timescales of neural dynamics on behavior.

## Main

Although brain activity exhibits fast temporal dynamics, whether these dynamics causally drive perception or cognition remains largely unresolved. Many theories posit that fine timescale activity controls communication within and between brains area^4,8–14^ which may govern sensory processing^9,15–18^, but also high level phenomenon such as attention^19^, social interaction^20^, movement planning^21^, memory^22^, and even conscious experience^23^. Yet causally probing the role of temporal dynamics on the millisecond timescale requires a technique to perturb these dynamics without altering firing rates (i.e., on the tens to hundreds of milliseconds timescale). Optogenetics^7,14,24,25^, electrical stimulation^26^, and pharmacological techniques^18^ can perturb network dynamics on fast timescales, but they also alter mean levels of neural activity, thus rendering them inadequate for separating the contribution of fast dynamics from slower, time-averaged activity on perception and behavior. It is possible to microstimulate or optogenetically stimulate at different frequencies or phase relationships to ongoing signals and compare the resulting behavioral effects^15,22,27,28^, but this still will not avoid a change in overall spike rates. For example, rhythmic stimulation of GABAergic neurons can potently entrain gamma oscillations but will also suppress spike rates through direct inhibition^15,27^. To overcome these issues, we sought to develop a new method that could alter fast neural dynamics while leaving average firing rates largely unchanged.

### Balanced bidirectional optogenetics

We hypothesized that a precise, bidirectional optogenetic approach could be used to alter neural dynamics while mitigating any changes on spike rate. The concept is broadly analogous to the ‘dynamic clamp’^2,30–33^, a single-cell method that can precisely control a neuron’s dynamics on specific timescales by injecting time varying membrane conductances. In both theory and experiment, introducing balanced input barrages of synaptic excitation and inhibition with specific temporal structure can independently regulate fast temporal dynamics and mean activity^34^. It has thus proven invaluable for testing theories of network-driven dynamics in isolated single neurons. If one could extend the precise control of the dynamic clamp to many neurons in a brain circuit simultaneously, one could implement the causal tests needed to distinguish the causal impact of different timescales of neural dynamics on perception and behavior. We thus sought to develop an approach with which we could simultaneously inject well-regulated and time varying excitation and inhibition into many neurons at a time.

To achieve this, we employed the bidirectional optogenetic modulator, BiPOLES^35^, that provides independent excitation and inhibition with different wavelengths of light. BiPOLES is a fusion of the fast, red light-sensitive optogenetic actuator, Chrimson^36^, and the fast, blue-light sensitive optogenetic silencer, GtACR2^37–39^, thus driving stoichiometric expression of an excitatory and inhibitory opsin that are also both potent and have fast kinetics (Fig. 1a). We speculated that this tool, when calibrated, could optically simulate precisely controlled excitatory and inhibitory conductances. By illuminating cells simultaneously with a specifically chosen ratio of blue and red light we might be able to approximate a ‘balanced’ state which should have a limited effect on mean spike rates. We could then exploit the fast kinetics of Chrimson and GtACR2 to dynamically inject asynchronous, but balanced barrages of excitation and inhibition which could overwrite endogenous cortical temporal dynamics and replace them with random structure.

**Figure 1:**
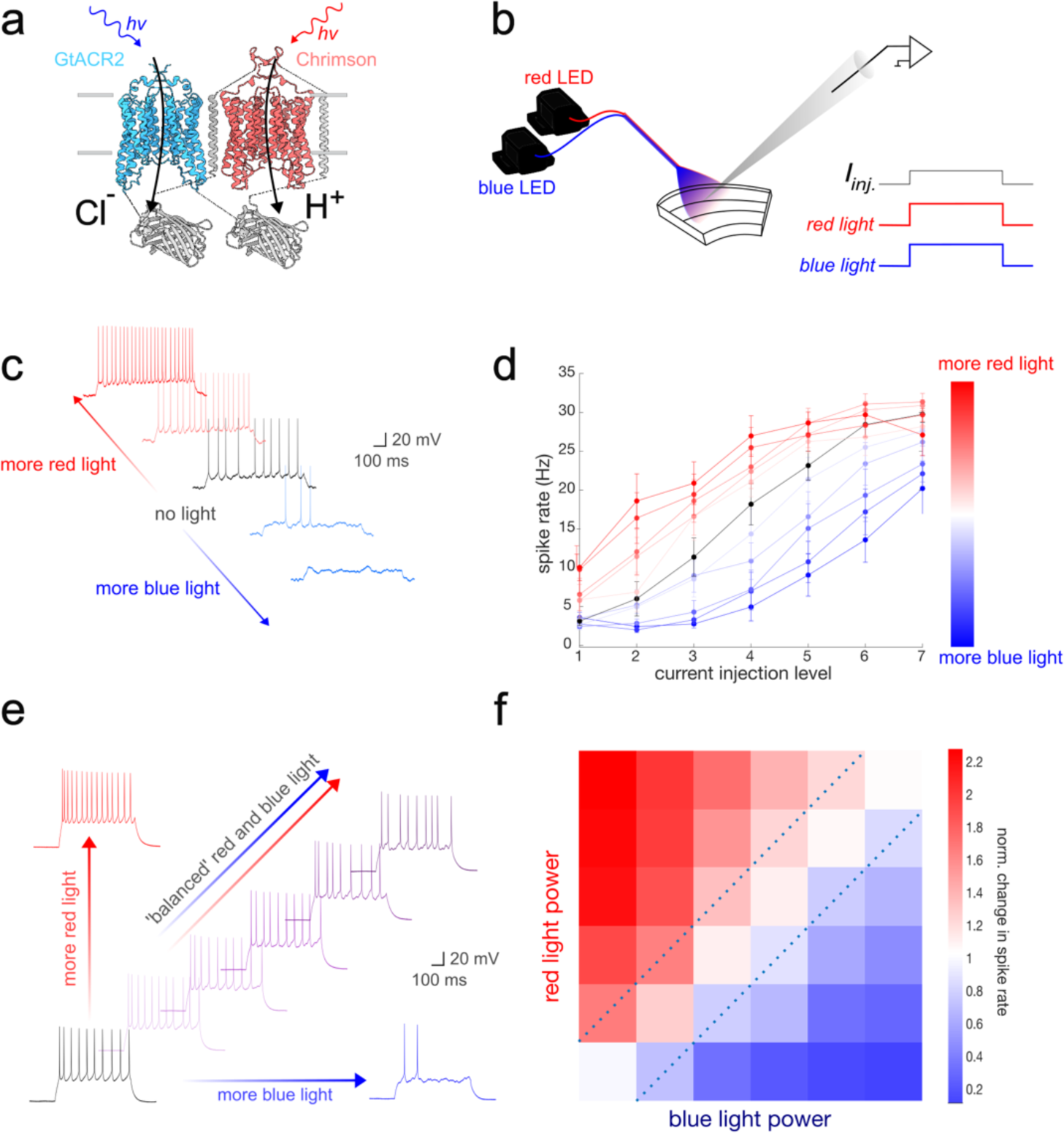
Balanced bidirectional optogenetic modulation. **a)** 3D schematic model of the bidirectional optogenetic construct BiPOLES, which is composed of the blue-light sensitive anion channel, GtACR2, fused to the red-light sensitive cation channel, Chrimson. **b)** Schematic of the recoding setup. A single cortical neuron in a brain slice is recorded intracellularly and illuminated with an optic fiber coupled simultaneously to a red and blue LED. Direct current injection through the electrode generates spiking activity, and waveforms of red and blue light modulate the neuron’s response. **c)** Example membrane potential traces of a cortical neuron showing positive and negative modulation of its firing rate with red and blue light, respectively. **d)** Average plot (n=6) of activation or suppression of cortical neuron firing rates with increasing levels of either red or blue light compared to no illumination (black). Error bars are s.e.m. **e)** Example membrane potential traces of a cortical neuron showing the modulation with red or blue light alone (red and blue, respectively) and with the ‘balanced’ red:blue light ratio (purple) of increasing total light power along the rising diagonal. **f)** Average plot (n=6) of bidirectional modulation of cortical neuron firing rates (normalized) across 36 different combinations of blue and red light intensities. The dashed diagonal lines outline the approximately balanced set of ratios that drive limited changes in firing rate.

To test these ideas, we first made intracellular recordings from cortical pyramidal cells in brain slices expressing a soma-targeted version of BiPOLES (Extended Figure 1a) and drove them to spike with direct current injection through the patch electrode while illuminating with different intensities of red and blue light (Fig. 1b). Conventional step pulses of either blue or red light could precisely increase or decrease the recorded neuron’s firing rate (Fig. 1c,d). To test if we could find a ‘balanced’ ratio of blue and red light that had a minimal impact on firing rate, we presented a range of ratios of blue to red light intensities (Fig. 1e). Indeed, we could find a narrow range of ratios that drove minimal net changes in mean activity (Fig. 1f, mean ratio: 0.33±0.05 blue to red, n=6, Extended Data Fig. 1b), presumably because the opposing conductances from Chrimson and GtACR2, combined with the net increase in the membrane shunt by both opsins, operationally canceled at the firing rate level. This demonstrates that simultaneously illuminating BiPOLES-expressing neurons with ‘balanced’ red and blue can cause no change in spike rates.

Next, we asked if injecting time varying, random barrages of excitation and inhibition (‘noise’), mimicking what naturally occurs *in vivo*^37–39^, could ablate fine temporal structure of neural activity. First, we assessed the conductances that noisy blue and red light would inject into BiPOLES-expressing neurons using voltage clamp. The light-induced membrane conductance for both wavelengths generate temporally matched waveforms, with some filtering (Extended Data Fig 1b, most likely a combination of the intrinsic kinetics of the opsins and some residual filtering through the patch clamp). Spectral analysis of the optogenetically injected conductances demonstrated that it could follow the light waveforms across a broad range of frequencies (Extended Data Fig. 1c). This demonstrates that the kinetics of Chrimson and GtACR2 are sufficient to control membrane conductance on fast timescales.

Next, we asked whether optogenetic injection of noisy excitation and inhibition could alter spike time. First, we first directly injected excitatory current into the patched neuron (via the patch pipette) either with frozen noise (i.e., a fixed, dynamic current injection that was precisely repeated trial to trial) or a sinusoidal current injection at 30Hz to drive them to spike in a quasi-physiological manner (Fig. 2a,c). Frozen noise drove highly repeatable time-locked sequences of action potentials in the recorded neurons, consistent with prior work^16,40–43^ (Fig. 2b, left). Similarly, sinusoidal current injection drove highly time-locked spiking activity (Fig. 2d, left). However, when we superimposed time-varying, asynchronous and noisy blue and red illumination profiles at the approximately ‘balanced’ ratio, we found that we could strongly interfere with the underlying temporal dynamics. Balanced red and blue illumination suppressed precise spike timing both for millisecond timescale locking of spikes to the frozen noise and for the ∼30 Hz spiking to the sinusoidal current injection (Fig. 2b,d,e, n = 21, p <0.005, Wilcoxon signed rank test). Importantly, even while this bidirectional optogenetic stimulation dramatically altered the temporal dynamics of neural spiking, it drove no significant change in average firing rates (Fig. 2f, n = 21, p=0.18, Wilcoxon signed rank test). These data demonstrate the feasibility for applying this approach *in vivo* to alter the temporal dynamics of endogenous sensory driven activity in the cortex.

**Figure 2:**
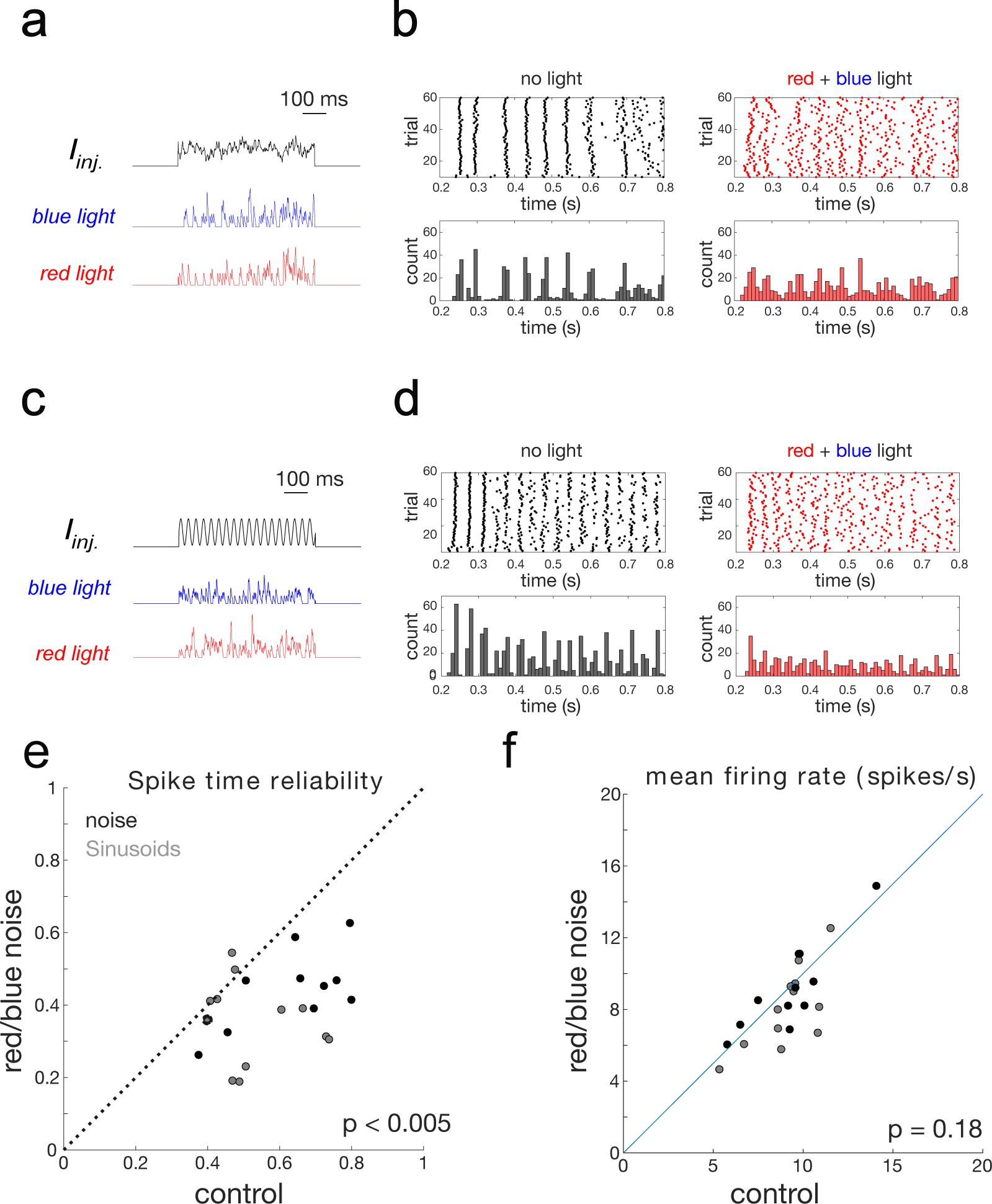
Randomizing fine timescale neural dynamics with balanced bidirectional optogenetic modulation. **a)** Example noisy current injection (black) and optogenetic illumination (red and blue) waveforms. The current injection waveform is identical across all trials, while the red and blue light waveforms change on every trial, but their mean intensities are fixed. **b)** Example raster plots (top) and spike histograms (bottom) of a cortical neuron in a brain slice in control conditions left (current injection only, no light) and with simultaneously red/blue light illumination. **c,d)** As in a,b) but for an oscillatory (30 Hz) current injection. **e)** Scatter plot of the spike time reliability (see Methods for analysis) comparing control (no light) and the red/blue light condition, combined across noisy and oscillatory current injection. N = 23 neurons, p <0.005, Wilcoxon signed rank test. Black points are for noisy current injection and gray points are for sinusoidal current injection. **f)** As in e) but for mean spike rate. N = 23 neurons, p = 0.18, Wilcoxon signed rank test.

### Balanced bidirectional optogenetics can alter fine timescale neural activity *in vivo*

To test this idea *in vivo*, we recorded primary visual (V1) cortical neurons juxtacelullarly in awake, head-fixed mice expressing BiPOLES broadly in the primary visual cortex (Extended Figure 1a). We drove physiological cortical activity with a high contrast visual stimuli while measuring and manipulating neural responses (Fig. 3a). We compared how various ratios of blue and red light would impact trial-averaged firing rates and fast cortical dynamics, hypothesizing that balanced square pulses of light would not affect either metric, while noisy light pulses would alter dynamics but not mean rates. Blue light alone suppressed the recorded neurons’ activity, while red light elevated spike rates (Fig. 3b, n = 20 units) confirming the basic efficacy of bidirectional manipulation *in vivo*. However, when we illuminated the cortex with a combination of red and blue light at various ratios, we could strongly mitigate any significant changes in visually evoked, trial-averaged firing rates whether using square pulses, noisy traces, or 30 Hz sinusoids (Fig. 3bm Extended Data Fig. 1c). Thus, even while balanced bidirectional optogenetic manipulation strongly increases total membrane conductance, at the firing rate level the excitation and inhibition operationally cancel.

**Figure 3:**
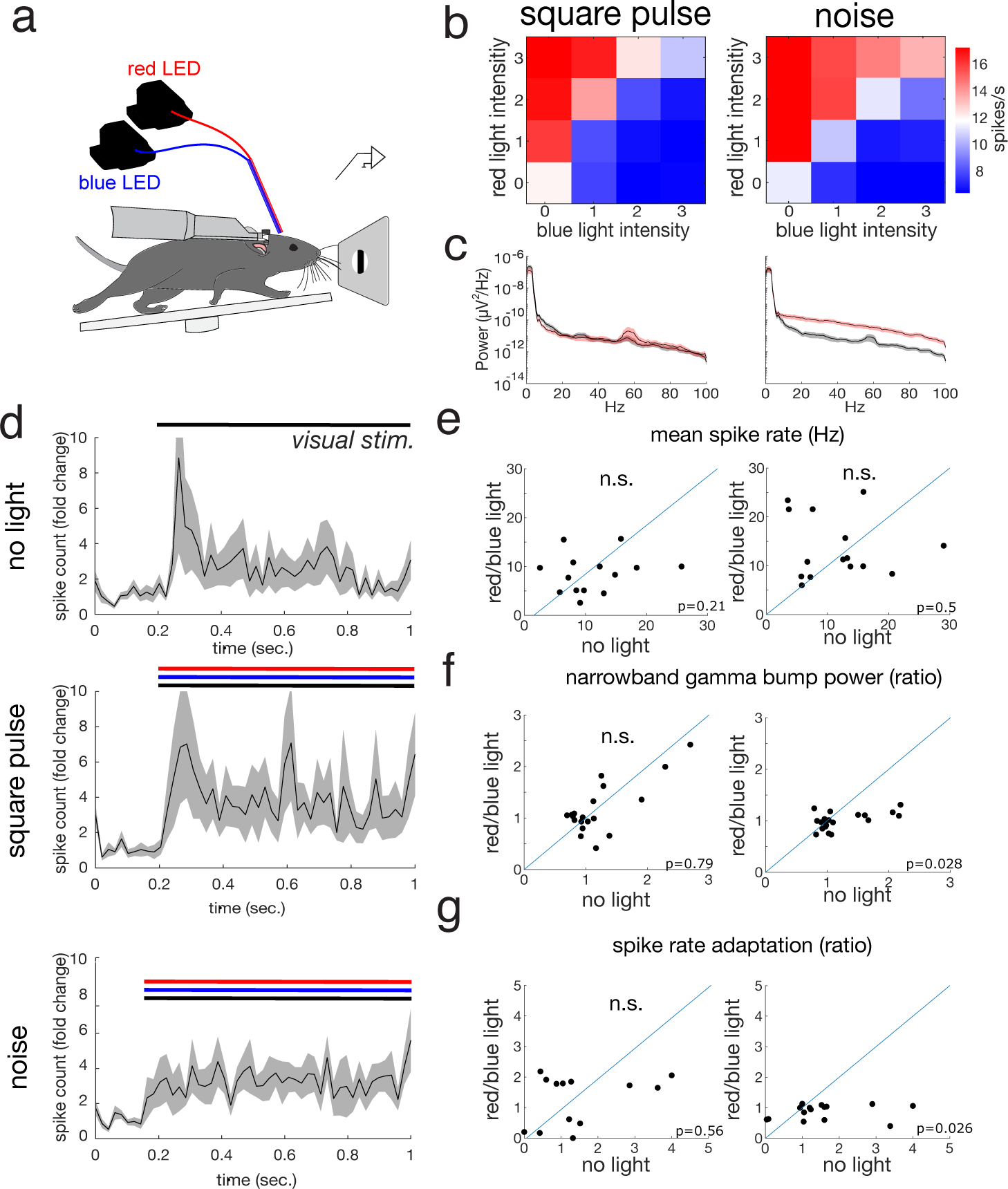
Randomizing fine timescale neural dynamics *in vivo* during visual stimulation. **a)** Schematic of the recording configuration. A head-fixed mouse runs on a circular treadmill while viewing a visual stimulus (grating) on a monitor. Stimulus location and orientation were matched to the unit’s preference. A single glass extracellular electrode simultaneously records the local field potential and a unit in the juxtacellular configuration. Blue and red LEDs are coupled to the same fiber which illuminate the recording site. **b)** Plots of average spike rates across all 48 conditions in each experiment (4 blue light intensities, 4 red light intensities in two temporal conditions: square pulses (left) and noisy waveforms (right); n=20 recordings. **c)** Average power spectral density plots (PSD) for the LFP recorded in two different conditions: square pulses (left) and noise (right). The PSD without light is plotted in gray, and with light (corresponding to the max red/light condition – upper right pixel of the plots in b), is plotted in red. Shaded area corresponds to s.e.m. **d)** Visually driven PSTH for the control (no light condition, top), and for square pulse (middle) and noisy light conditions (bottom). Errors bars are s.e.m. e) Scatter plot of mean spike rate for the square pulse (left) and noise condition (right) between control (no light) and max red/blue light. **f)** Scatter plot of the narrowband gamma ‘bump’ power ratio in the PSD for the square pulse (left) and noise condition (rigt) between control (no light) and max red/blue light. For analysis see Methods. **g)** Scatter plot of the spike rate adaptation ratio in the PSTH for the square pulse (left) and noise condition (right) between control (no light) and max red/blue light.

Next, we asked how these different temporal patterns of balanced illumination impacted endogenous temporal dynamics. To quantify changes in temporal structure we first analyzed the effect on the power spectrum of the cortical local field potential (LFP) as a holistic metric of neural dynamics. In control conditions (no optogenetic illumination) with high contrast visual stimulation, the LFP power spectrum showed broad power with an extra ‘bump’ in the gamma band centered on ∼57 Hz, consistent with prior findings^44–47^ (Fig. 3c). Illuminating with balanced, square pulses of red and blue light (corresponding to the upper right pixel of the plots in Fig. 3b) did not dramatically alter the power spectrum (Fig. 3c, left) nor did it affect mean spike rate (Fig. 3d,e, n = 14 units, p=0.21, Wilcoxon signed rank test). Illuminating with the same average red and blue light power, but using noisy illumination profiles, also didn’t affect mean spike rate (Fig. 3d,f, n = 14 units, p=0.50, Wilcoxon signed rank test), but substantially restructured the power spectrum, increasing power across a spectral band (Fig. 3c, center). It also suppressed the ∼57 Hz gamma band ‘bump’ (Fig. 3f, n = 20 recordings, p=0.028, Wilcoxon signed rank test), unlike the square pulses which had no significant effect on it (Fig. 3e, n = 20 recordings, p=0.79, Wilcoxon signed rank test). We quantitively compared the overall effects of balanced square pulse and noise illumination on the LFP by computing a similarity index of the power spectrum in control conditions as compared to balanced illumination. Noise stimulation cause a dramatically greater drop in similarity than did square pulse stimulation (Extended Data Fig 1b, n = 20 recordings, p=0.0003, Wilcoxon signed rank test), demonstrating that balanced optogenetic noise injection was far more effective at restructuring the overall temporal dynamics of the local network. To test if we could elevate specific frequency bands, we found that illuminating with balanced 30 Hz sinusoids of blue and red light potently increased power in the 30 Hz and its harmonics, even while it didn’t change spike rate (Extended Data Fig. 1c-g).

Lastly, we examined the temporal dynamics in the peri-stimulus time histograms (PSTHs) of the visually evoked response of the juxtacellularly recording units. In control conditions, the units showed a fast and strong response to the visual stimulus, peaking ∼60 ms after visual stimulus onset, and then relaxing to a more sustained response for the duration of the stimulus (Fig. 3d, top). This fast and adapting response was only a feature of high contrast stimuli, but largely absent at low contrasts (Extended Data fig. 1h). When simultaneously illuminating with balanced square pulses of red and blue light, the shape of the PSTH was largely unaltered, showing no change in this peak/sustained spike rate (Fig. 3d,g, n = 13, p=0.56, Wilcoxon signed rank test). However, noise stimulation largely eliminated the early fast peak of the visual response (Fig. 3d,g, n = 15 units, p=0.026, Wilcoxon signed rank test), while 30 Hz sinusoids drove an oscillatory profile in the PSTH (Extended Data Fig 1e). Taken together, these electrophysiological data demonstrate that bidirectional optogenetic control with BiPOLES *in vivo* provides a means to substantially alter the temporal structure of physiological cortical activity while avoiding significant changes in firing rates.

### Altering neural dynamics suppresses visually guided behavior

We next took advantage of this bidirectional optogenetic system to test whether altering temporal dynamics in the primary visual cortex (V1) would influence sensory perception. We trained mice on a contrast detection task in which they had to lick to the presentation of a drifting grating of various contrasts^1,3,5,9,22,48–50^ (Fig. 4a). Illuminating V1 in BiPOLES-expressing mice with long step pulses of blue light alone, which potently silences cortical activity (see above), dramatically reduced performance on the task, manifesting as a rightward shift in the psychometric function and an increase in detection threshold (Fig. 4b, Extended Data Fig. 3a, detection thresholds, light off: 5.0 ± 0.5%, blue light: 21.8 ± 5.0%, p=0.001 Tukey post-hoc test). This result is consistent with prior findings that silencing the cortex reduces contrast sensitivity^44,47,48^. Conversely, illuminating BiPOLES-expressing mice with red light alone resulted in a potent increase the animal’s response rate at low contrasts, consistent with the notion that elevating V1 activity is sufficient to trigger perceptual decisions (Fig. 4b, Extended Data Fig 3a, detection thresholds, light off: 5.0 ± 0.5%, red light: 9.4 ± 1.5%, p=0.94 Tukey post-hoc test). Illuminating mice that did not express BiPOLES with blue or red light had no effect on the animals’ psychometric functions (n=3, Extended Data Fig. 4b, p=0.27 rmANOVA).

**Figure 4:**
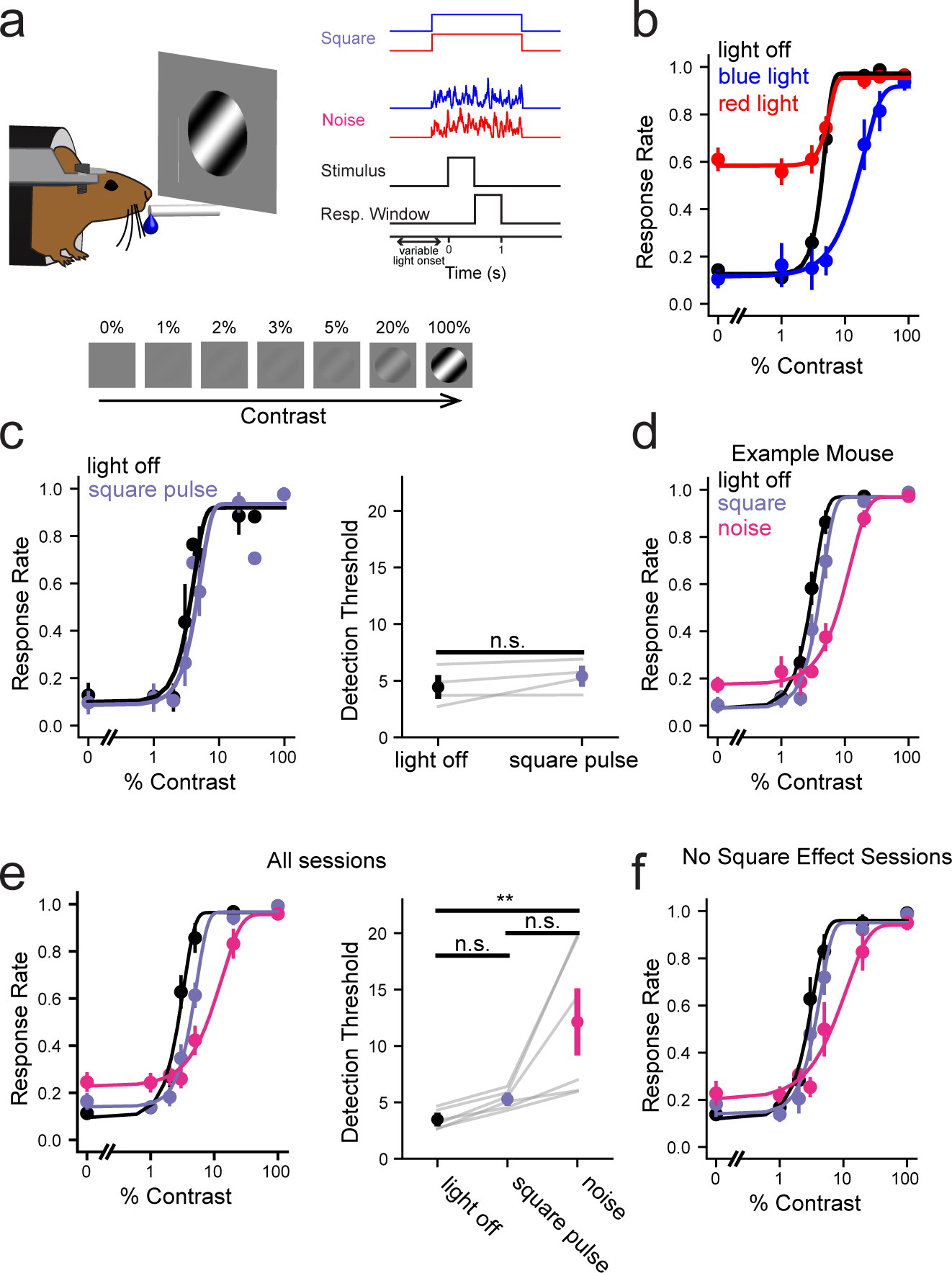
**a)** Left: schematic of the behavioral setup. Right: time course of behavioral trials and different illumination conditions (right). Bottom: schematic of the visual stimuli used. **b)** Average psychometric curves in experiments where blue light only or red light only was delivered to visual cortex on a subset of trials. n=6 mice. p=0.0015 rmANOVA, Tukey post-hoc tests: no light vs red: p=0.94, no light vs blue: p=0.001, blue vs red: p=0.002. **c)** Left, as in b) but for sessions in which square pulses of blue and red light were delivered simultaneously at the operationally balanced level. Right, average detection thresholds (% Contrast) for light off and square pulse conditions. Light gray lines, individual mice. p=0.17, paired t-test, n=4 mice. **d)** Psychometric curves for an example mouse in which no light (black) trials, square pulses of red and blue light (purple) and noisy red and blue light waveforms (magenta) were delivered to V1. **e)** Left: As in d) but for average performance over n=6 mice (10-18 sessions per mouse). Right: detection thresholds for each mouse in each condition. Light gray lines, individual mice. p=0.0033, repeated measures ANOVA, Tukey post-hoc tests: noise vs no light, p=0.004, noise vs square: p=0.035, none vs square: p=0.8. **f)**: As in e) but for sessions in which square pulses of red and blue light was determined to have no effect via within-session bootstrapping of the hit rate at middle contrasts (p=0.0060 repeated measures ANOVA. Tukey post-hoc tests: noise vs no light: p=0.0099, noise vs square: p=0.0349, square vs no light: p=0.8003).

Next, we asked if we could ‘balance’ the effect of blue light suppression by simultaneously illuminating with red light. Over several behavioral sessions we tested increasingly higher intensities of red light until we found red light levels that shifted the psychometric leftwards such that it would be indistinguishable from control (no light) conditions (Fig. 4c, Extended Data Fig. 4c, p=0.17 paired t-test). These data demonstrate that we could use bidirectional optogenetic control with conventional step pulses of light to identify a blue to red light intensity ratio that would result in minimal effects on behavior. Without direct measurement of all V1 neurons’ spike rates in these animals we cannot be sure whether there was a net change in the firing rate of at least some V1 neurons. However, since blue light on its own suppresses performance, and red light on its own increases response rate at low contrast, the operationally ‘balanced’ ratio of red to blue light we used here is unlikely to have caused a substantial change in mean V1 activity.

Equipped with this ratio, we next used it to test the causal impact of endogenous temporal structure on sensory perception. To do so, we interleaved three trial types in single sessions: no light (‘control’), simultaneous square step pulses of red and blue in the ‘balanced’ condition (‘square pulse’), and noisy profiles of blue and red light (‘noise’). Importantly, the mean intensity of both the red and blue light in the noise condition (i.e., total illumination power across the trial) was identical to the step pulse condition. To avoid cueing the mouse to any trial information, all LED conditions started at a random time prior to stimulus onset (Methods). As we showed above, noisy, balanced, bidirectional optogenetic stimulation, but not square pulses, dramatically alters the temporal structure of neural activity while driving limited changes in total firing rates. If the natural temporal dynamics of cortical activity is critical for sensory perception, then the mice should perform significantly worse during noise stimulation than on the other two conditions. Conversely, if the endogenous temporal structure is not needed for behavioral performance, there should be no difference between noise and square pulse stimulation. Consistent with the latter hypothesis, mice performed substantially worse during noise stimulation than for square pulses (Fig. 4d.e), even though the mean level of the light illumination was identical between the two conditions. When averaged across all such sessions, square pulse stimulation caused no significant change in the animals’ detection threshold, but noise stimulation substantially reduced it (Fig. 4e p=0.0033, rmANOVA, Tukey post-hoc tests: noise vs no light, p=0.004, noise vs square: p=0.035, none vs square: p=0.8.).

It is possible that that ratio of blue and red light we used in these behavioral experiments might still have elevated or suppressed mean firing rates in V1. If it had driven a substantial enhancement, it should have augmented the response rate at low contrasts, which is not what we observed. Conversely, if it had driven a substantial suppression, both square pulses and noisy optogenetic stimulation should have reduce performance across contrasts. Indeed, in 36/77 sessions we observed a significant, albeit slight negative effect of square pulse stimulation on performance, suggesting that the ratio we used may have not been exactly balanced. Thus, as a stringent test of the hypothesis that spike timing causally influences perception, we excluded any sessions where the square pulses of red and blue light caused a detectable effect on performance. Even in this subset of sessions (41/77) noisy bidirectional optogenetic stimulation caused a substantial reduction in stimulus detection (Fig. 4f), arguing that the alteration of spike timing *per se* is what drove the reduction in task performance.

The methodology we used here alters neural dynamics on both fast (∼milliseconds) and medium (tens of milliseconds) timescales, without altering it on longer time scales (∼seconds). Thus, it could be possible that the behavioral impact of noise stimulation on contrast detection could be ascribed to specifically suppressing the fast peak of neural activity that occurs in the first ∼100 milliseconds of the visual response, even while it leaves the total number of visually driven spikes unaltered. However, the effect of noisy optogenetic stimulation on performance manifested only at low contrasts, and low contrasts do not drive this fast, adapting visual response, largely ruling out this possible explanation (Extended Data Fig. 3d,e; n = 22 mice). To further test this idea, we silenced V1 activity only during the initial transient of the visual response (∼60 ms of the visual response) and compared its effect to silencing during the entire visual stimulus and response period as done above. We found that such silencing had only a minimal effect on behavior (Extended Data Fig. 3f,g, n = 3 mice), ruling out any effect of the bidirectional noise stimulus on the initial fast visual response transient as a major explanation of its greater impact on behavioral performance. Thus, by dissociating the effect of fine timescale neural dynamics from slower changes in firing rates using bidirectional optogenetic modulation, these experiments show that endogenous, fast temporal dynamics in the cortex are necessary for sensory perception.

### Conclusions

Accumulating evidence over several decades has argued both for and against the notion that the endogenously driven fine temporal dynamics of cortical activity causally contributes to neural computation and perception^1,3,5,9,22,49–51^. On one hand, sensory stimulation frequently drives potent temporal correlations in neural activity within and across cortical areas whose strength and frequency correlates with sensory detection and attention^51^. Conversely, other studies have argued that this fast temporal structure can be absent for natural stimuli^49^, inconsistent for multiple stimuli of varying contrast^52^, or simply unrelated to perceptual output^52^. Recently, multiple studies have shown that optogenetic modulation of neural synchrony or spike timing can influence behavior^15,22,27,53^, supporting the timing hypothesis. However, because these studies employed unidirectional optogenetic stimulation of excitatory or inhibitory neurons they also altered mean firing rates, thus leaving it potentially uncertain as to whether the behavioral change was purely due to alterations in temporal dynamics or could also be ascribed to alterations in mean activity.

By developing and leveraging a novel bidirectional optogenetic approach that uncouples perturbations to spike timing from perturbations to spike rates, we executed a direct causal test of fast temporal dynamics in sensory perception. Our results show that randomizing the endogenous temporal structure in mouse V1 potently suppresses performance on a sensory detection task. This argues that activity on fine timescales in the cortex is central to neural computation and behavior. Notably, although we could randomize the temporal dynamics of the V1 population, our wide field illumination is likely to have generated strong synchrony in the illuminated population on each trial. The timing of this synchronization was random on each trial and thus had no consistent reference to the endogenous network structure nor to the behavior. Since spike timing synchrony has been proposed to facilitate the propagation of activity^11^, this artificial increase in synchrony within each trial could have facilitated sensory perception, but this is not what we observed.

Another possibility is that noisy, balanced bidirectional optogenetic stimulation could have impacted behavior by increasing correlated variability in V1 (i.e., noise or ‘spike count’ correlations). That is, even while it would not change mean spike rates across all trials, it would lead to increased variability across trials, and this variability would likely be correlated across neurons since they receive the identical noisy balanced optogenetic input on each trial. Increased correlated variability can degrade performance in sensory tasks because it limits the advantages of signal detection gained from averaging across neurons. Thus, it could be possible that noisy bidirectional optogenetic stimulation degraded behavior due to its potential impact on spike count correlations, not on fast temporal dynamics. However, since adding spikes on a given trial should increase the probability of detection, while removing spikes should decrease it, these effects should have canceled in the average performance of the mouse, but they did not: noisy bidirectional optogenetic modulation strongly suppressed performance. Thus, we instead conclude that the altered temporal pattern of activity interfered with endogenous temporal dynamics to alter perception, and that any increase in correlated activity is unlikely have explained the behavioral effects.

Future experiments could use this approach to determine which class or spectral band of fast dynamics is most beneficial for sensory perception. This could be done by elevating or suppressing rhythmic synchrony within or between brain areas in specific frequency bands (e.g., alpha, beta, ‘low gamma’ or ‘high gamma’) without directly altering mean rate, while assessing the effects on behavior. Similarly, for addressing whether specific spike sequences are critical, closed-looped perturbations which trigger optogenetic stimulation when spike sequences occur^22^, could be used to test their role in perception and behavior. Furthermore, it is possible that spike timing is important for certain periods within a behavior (i.e., a sensory detection trial) but not others. This could be addressed by disorganizing neural activity (without changing mean rate) in specific periods of a behavioral trial but not others. Moreover, any of these dynamical perturbation approaches can be targeted to specific transcriptionally or projection-target defined cell types by expressing BiPOLES in driver lines or with a retrograde virus. Doing so would allow cell-type specific causal dissection of timing codes in brain circuits. Relatedly, it could be beneficial to implement the bidirectional optogenetic approach we developed with high resolution two photon excitation. This would allow investigators not only to target specific cell types, but even to target specific, functionally defined ensembles of cells using holographic illumination.

### Experimental Methods

All experiments were performed with the authorization of UC Berkeley’s Animal Care and Use Committee.

#### Animals and viral injections

All mice utilized in the study were C57/B6 wild type mice, obtained either from Charles River or bred in-house. Both male and female mice were utilized equally in the experiments. For viral injection, neonatal mice aged between P3-P5 were cryoanesthetized and secured with tape in a ceramic mold. Intracranial injections were administered using a NanoJect II system (Drummond) equipped with a beveled glass injection needle (WPI), loaded with 0.5-1 μL of AAV9-CaMK2II-somBiPOLES-mCerulean (Addgene Plasmid #154948). The injection sites and needle insertions were controlled using a micromanipulator (Sutter, MP-285). Three injection sites were targeted within V1, positioned relative to the lambda suture at 0.0 mm AP and 2.1 mm lateral left, at four different depths (ranging from 100-400 μm). 27nL of solution was delivered per location.

#### Headpost surgery and craniotomy

All procedures were conducted in compliance with the guidelines by the Animal Care and Use Committee of the University of California, Berkeley. A custom stainless-steel headplate was surgically implanted for headpost implantation. Adult mice aged between P51 and P159 were anesthetized with 2–3% isoflurane and positioned in a stereotaxic apparatus. The scalp was removed and the skull lightly scored using a drill bit, then vetbond was applied to the skull surface. The headplate was positioned over V1 (V1; 2.7mm lateral, 0mm posterior to lambda) and attached with dental cement (Metabond) and the remaining exposed skull was covered with Metabond. For optical access to V1 in behavioral experiments (Figure 4), a cranial window was implanted prior to covering the skull with dental cement. A 3.5mm biopsy bunch was used to remove the skull and a cranial window consisting of two, 3-mm and one 5-mm coverslip, was placed and secured with Metabond (C&B). By the time of optogenetic behavioral experiments the dura under the cranial window had thickened and become semi-transparent. Following the procedure, mice were placed in a heated recovery cage to facilitate recuperation. Mice received buprenorphine and meloxicam for postoperative pain management and were allowed a minimum of three days to recover before undergoing water restriction for experimental procedures.

#### Brain slice electrophysiology and optogenetics

Injected mice P14-P28 were deeply anesthetized with isoflurane and decapitated. The injected (left) hemisphere was sectioned with a Microslicer Zero 1N (Ted Pella) in modified sucrose enriched ACSF, briefly kept at 32 degrees and then at room temperature. Slices were stored in the same buffer until use. Whole cell patch clamp recording under visual and epifluorescence guidance were performed with a Scientifica slice scope equipped Sutter micromanipulators (MP-285), a Molecular Devices 700B Multiclamp amplifier, and a Lumencor Spectra X for epifluorescence illumination. All data acquisition and instrumentation were controlled with custom code written in Matlab. Optogenetic illumination was performed with two fiber coupled LEDs (Thorlabs) with center wavelengths at 455 and 617 nm and merged with a 400-micron core 2˗2 fiber optic coupler (Thorlabs TT400R5S2B). The tip of one of the outputs (after removal of any termination and cladding) was mounted on a micromanipulator and position underneath the microscope object <5mm from the patched cell. Output power per color, measured at the tip of the fiber, measured 0.1-1.5 mW. Maximum power was slightly adjusted between slices to adjust for variations in opsin expression level. Single cortical pyramidal cells (L2/3 or L5) were patched with low resistance borosilicate patch pipettes (Sutter instruments), filled with a potassium based internal solution (290 mOSM). The extracellular solution was contained ACSF with 2.5 mM calcium and 1mM Kynurenic acid to block all glutamatergic synaptic transmission and warmed to 32 degrees Celsius. The light waveforms were generated via voltage command signals from the National Instruments data acquisition device.

#### Experiment in **Fig. 1c,d**

Patched pyramidal cells were injected with seven levels of a current to generate a range of firing rates. For each current level random zero mean noise was added to mimic naturalistic synaptic noise. For each cell the range of current levels was adjusted to maximize the dynamic range of the cell’s spike output. On top of these current injections, various levels of red and blue light, separately and without any added noise, were used to test for increases or decreases in mean firing rates. Spikes were automatically detected via a 0 mV threshold.

#### Experiment in **Fig. 1e,f**

Patched pyramidal cells (a different group than for the experiment in Fig. 1d) were injected with a single level of direct current to evoke a mean spike rate of ∼10 Hz. On top of this current injection, six levels of red and six levels of blue light (including a zero power level for each color) were projected onto the slice in all 36 combinations. The illumination time series were generated as follows: two random spike trains with Poisson statistics were generated for each trial, one for red light and one for blue light. Each spike in the spike train was convolved with an alpha function designed to simulate single excitatory and inhibitory postsynaptic conductances. This process generated random, noisy, and uncorrelated blue and red light illumination time series. To generate the 6 different mean levels of red or blue light, these traces were multiplicatively scaled with 6 preset gains for each color. All trial types were randomly interleaved. Spikes were automatically detected via a 0 mV threshold.

#### Experiment in **Fig. 2**

Patched pyramidal cells were injected with a single, non-varying (‘frozen’) noise trace generated by convolving a random Poisson spike train with an alpha function or they were injected with a 30Hz rectified sinusoid. The amplitude of either the noise trace or the sinusoid was scaled for each neuron to evoked ∼5-10 Hz mean firing rate. On top of this current injection a single combination of noisy red and blue light times series (generated as above) were projected via the optic fiber. All trial types were randomly interleaved. Spikes were automatically detected via a 0 mV threshold. To quantify spike time reliability, we employed previously used method^54^ which computes the mean correlation of spike times across all pairs of trials. In Fig. 2e,f data is collated across the noise and 30Hz current injections because in both conditions there was a significant effect on spike time reliability and no significant effect on mean spike rate. For the conductance recordings in Extended Data Fig. 1c,d, BiPOLES-expressing pyramidal neurons were patched in the voltage clamp mode with an internal where cesium replaced potassium, and included QX-314 and TEA. Chrimson currents were measured at -70mV when illuminating with red light only, and GtACR2 currents were measured at 0mV when illuminating with blue light only.

#### In vivo juxtacellular electrophysiology and optogenetics

Adult mice (P35-P71) were anesthetized and implanted with a stainless steel headplate for fixation. The location of V1 was marked and the exposed skull was protected with a thin layer of dental acrylic (C&B Metabond). Mice were habituated to head fixation and locomotion on a 6” diameter circular treadmill over several days. On the day of the experiment, mice were briefly anesthetized with 2% isoflurane and a small <300 micron craniotomy was made by thinning the skull until it was nearly transparent with a drill bit and opening a small hole for electrode insertion with a 27 gauge needle. The dura was left intact. The mouse was fixed on the recording rig and allowed to recover from anesthesia for ∼15 minutes prior to recording. A single glass electrode was used to record LFP and well isolated units via juxtacellular recording. Glass electrodes were used because they avoid the confounding effect of light-induced photo-electric artifacts that are particularly problematic with fluctuating light waveforms when using silicon based multi-electrode arrays. The glass electrode (∼4 Megaohm) was filled with ACSF and lowered perpendicularly into V1 and moved with 2 micron steps. An Axopatch 200B set in I=0 mode, gain = 500 was used for extracellular recording and all data acquisition (NIDAQ) and instrument control was conducted via custom code written in Matlab. The same 2-color fiber coupled LED system was used for *in vivo* optogenetics and placed ∼3 mm from the craniotomy. Average maximum illumination power, measured at the tip of the fiber, was 0.63±0.9 mW for red light and 0.73±15 mW for blue light. The maximum light power was slightly adjusted in each experiment to normalize for changes in opsin expression level. During electrode lowering, brief pulses of red light were used to photo-stimulate cortical neurons (via Chrimson) to identify well isolated BiPOLES-expressing units. When such a unit was encountered the electrode was lowered several microns further to achieve the juxtacellular configuration, often yielding very well isolated spikes of that unit. Units were then probed with a series of drifting gratings (Pyschophysics toolbox) ∼20 degrees in diameter. The stimulus monitor was a gamma-corrected 23” Eizo LCD monitor positioned ∼15 cm from the mouse’s eye. The location and the orientation of the grating was adjusted to maximize the visually evoked firing rate of the isolated unit. Contrast was set to 50%. The experiment then commenced. On all trials (1 second length with a 2 second intertrial interval) the selected grating appeared 200 ms after trial onset for 800 ms and varied in contrast. The red and/or blue light turned on co-incident with the grating (except for 5 units where the light came on at the beginning of the trial) and at 16 different ratios of intensity (including the zero power level for each color) and across three temporal dynamic conditions: square pulses, noisy traces (generated as above), and a rectified 30 Hz sinusoid. All trial types were randomly interleaved. Twenty two juxtacellularly recorded units were recorded in five different mice, across seven different sessions, at depths spanning ∼236-705 microns below the pia.

##### Analysis

After high pass filtering, spikes were detected via a threshold set at 4.5-fold the standard deviation of the entire input trace. To compute power spectral density (PSD) the LFP was first downsampled to 200 Hz, and the PSD was estimated via the multitaper method (‘pmtm’ in Matlab R2021b). PSD similarity in each condition for each recording was computed via their cosine similarity. The estimate of narrowband gamma was computed by taking the ratio of the integral of the PSD in a 10 Hz band around the narrowband bump (typically 55 Hz) and dividing that by the sum of the integral in two adjacent 5 Hz side bands. The spike rate adaptation ratio was computed by measuring the ratio between the peak mean spike rate (measured 60-140 ms after the onset of the visual stimulus), to the sustained mean spike rate (measured 220-600 ms after the onset of the visual stimulus). For each analysis in Fig. 3, experiments with fewer than four trials in the maximum blue/red light power condition were excluded from spike rate analysis.

### Extracellular multi-electrode array recording

For data in Extended Data Fig. 3f,g, contrast responses were collected and analyzed following prior methodology^55^ using Neuronexus 16-channel silicon probes. The adaptation ratio of the PSTH was computed as the firing rate from 60-140 ms after visual stimulus onset divided by the firing rate bewteen 220 and 600 ms after visual stimulus onset.

### Water Restriction

For water restriction, each animal’s initial weight was recorded and then it was placed on a high fat diet ad libitum with restricted access to water. During training sessions, the mice received the majority of their water intake. Following training, the mice were weighed, and if their weight dropped below 75% of their initial weight, they were provided with additional water. The weight and overall health status of the mice, including fur and nail condition, gait, and posture, were monitored daily.

### Behavioral training

#### Apparatus

Two apparatuses were used for behavior. The first, used for training and for initial characterization of optogenetic effects (Figure 4a-c), was controlled real-time using an Arduino DUE synchronized with a Raspberry Pi 3+/4+, which interfaced with custom-written code in Python, Arduino, and Java (Bounds et al., 2021). Visual stimuli were generated with custom-written Python code and presented on a gamma-corrected LCD monitor (Podofo, 15.7 × 8.9 cm, 1024×600 pixels, 60 Hz refresh rate) positioned 12 cm from the right eye in a 30-degree angle from the longitudinal axis.

The second apparatus, used for Figure 4d was written in python using Psychopy^56^ **(citation)** and used a NIDAQ control box to control the LEDs and water valve. Stimuli were displayed on a gamma-corrected monitor (Adafruit 2406, 18 x 12 cm, 800 x 480 pixels).

In both cases, mouse licking was detected by changes in capacitive load using a capacitive touch sensor (Adafruit, AT42QT1070), creating a circuit between the 0.05-inch diameter steel lick port and the mouse’s tongue. Water delivery was achieved by gravity through the lick port using a 2-way normally closed isolation valve control (Neptune Research Inc.). The entire setup was mounted on an 8.0 × 8.0 × 0.5-inch aluminum breadboard (MB8; Thorlabs Inc.) and enclosed in a light isolation box (80/20).

#### Behavioral training and task design

Training commenced following surgery for headplate and cranial window implantation, as well as the subsequent recovery period (7 days) and water restriction (5–8 days). Before training, mice underwent two 30-minute habituation sessions to head fixation a tube with the monitor activated. The training process comprised four main stages. Initially, mice underwent classical conditioning to associate licking with a visual stimulus which consisted of a drifting sinusoidal grating square patch (2Hz, 0.08 cycles/degree, 600 ms, 100% contrast, 34 visual degrees, against a black background luminescence). Each trial presented this visual stimulus paired with a water reward at the onset of the response window. Catch trials were absent during this stage. Advancement to the operant conditioning phase occurred once mice consistently licked within the response window in over 80% of trials.

During the second training phase, mice learned to respond to visual stimuli, with 25% of trials being catch trials. To receive a water reward, mice had to lick within a specified window upon stimulus presentation while refraining from licking to avoid time-out penalties. Initial stimuli characteristics were consistent with stage one (100% contrast, 34 visual degrees, against a black background). Upon meeting performance criteria (>90% hit rate and <30% false alarm rate) for two consecutive days, mice progressed to stage three. In this stage, background luminance gradually increased to mean luminance, and stimulus size decreased to 18 visual degrees through session-to-session adjustments while maintaining performance criteria. After completing ramping, mice had to sustain performance for one day to advance. Stage four required mice to detect stimuli with varying contrasts. Optogenetic testing utilized six contrast levels, with catch trials representing 25% of all trials.

#### Optogenetic stimulation and noise pattern generation during behavior

Optogenetic stimulation during the contrast detection task utilized the same hardware as described above. The optic fiber was positioned via a micromanipulator above the V1 region at the rear of the window. Light intensity was calibrated at the fiber tip using a power meter (PM160T, Thorlabs Inc.). For Figure 4b-c, LEDs were controlled by Arduino. To test the bidirectional action of BiPOLES in behavior (Fig. 4b), LEDs were turned on simultaneously with the stimulus onset using 455 nm (4.0-5.0 mW) for silencing and 617 nm (0.05 mW) for activation. To determine the appropriate ratios to suppress individual opsin action, we conducted individual sessions with different blue-to-red ratios of 1:40, 1:30, 1:20, and 1:10 in four out of the six mice. In these sessions (Fig. 4b), LEDs were randomly activated between 300 and 3000 ms prior to stimulus onset until the end of the response window in 33% of trials. Once the appropriate ratio was determined, the mouse transitioned to the noise stimulation phase.

For noise stimulation experiments, data acquisition and generation of uncorrelated noise illumination patterns for optogenetic photostimulation was performed using a real-time with custom code written in Python using the National Instruments Data Acquisition (DAQ) Toolbox (NI). The noise patterns were generated in a random fashion for each light color and uncorrelated from each other, as described above. The appropriate mean light stimulation power was originally assessed in a pilot set of mice to identify a blue light power that effectively suppressed performance on its own, and then a red-light power that when combined with this blue light power minimally affected performance. In the final set of experiments, blue light power at the tip of the fiber was set to 2 – 3 mW and power for red light was set to one tenth of the blue light power in each mouse. The higher blue light powers needed for optogenetic suppression compared to the acute craniotomy experiments was likely explained by the thickening and loss of transparency of the dura (see above). Light was administered at a random time between 300 to 1500 ms prior to stimulus onset until the end of the response window. Square, noise, and LED off trials occurred as an equal percentage of trials. Due to the need for the nidaq control to finish a pattern prior to starting a new one, in some trials inter trial false alarms prevented the LED from being turned on at the correct timing, these trials were excluded from analysis for all trial types (including light off when no LED was shown). To prevent LED illumination *per se* from affecting behavioral performance, a custom-designed light-blocking cone was affixed to the mouse head, and blue LED masking lights were illuminated throughout the session. Additionally, to minimize extraneous cues, the left eye was covered with a plastic eye patch.

### Behavioral analysis

Sessions with hit rates at 100% contrast < 85% or with false alarm rates > 30% were excluded from analysis. For psychometric curves and calculation of detection threshold, behavioral data were fit with a modified Weibull function with false alarm and lapse rate allowed to vary.

Detection thresholds were the 63% point on the curve.

### Data and code availability

Electrophysiological and behavioral data and code are available on request.

## Acknowledgements

We thank Anushree Gupte for assistance with behavioral training, Corliss Lay for assistance with histology, K. Gopakumar, S. Sridharan, and J. Beyer for scientific and logistical support.

Additionally, we thank D. Feldman, M. Feller, and H. Shin for comments on the manuscript, and S. Brohawn for 3D schematic of BiPOLES. This work was funded by NEI grant R01EY023756 and RF1NS128772, and a fellowship from the Chan-Zuckerberg Biohub.

## Author contributions

H. A. conceived of the project. H.A performed all bidirectional electrophysiological experiments and analysis. D.Q. performed all viral injections and behavioral experiments. H.A.B. developed the behavioral task, supervised behavioral experiments, and conducted statistical analyses. J.V. collected and analyzed the visual contrast response functions. H.A. wrote the manuscript.

---

**Extended Data Figure 1:**
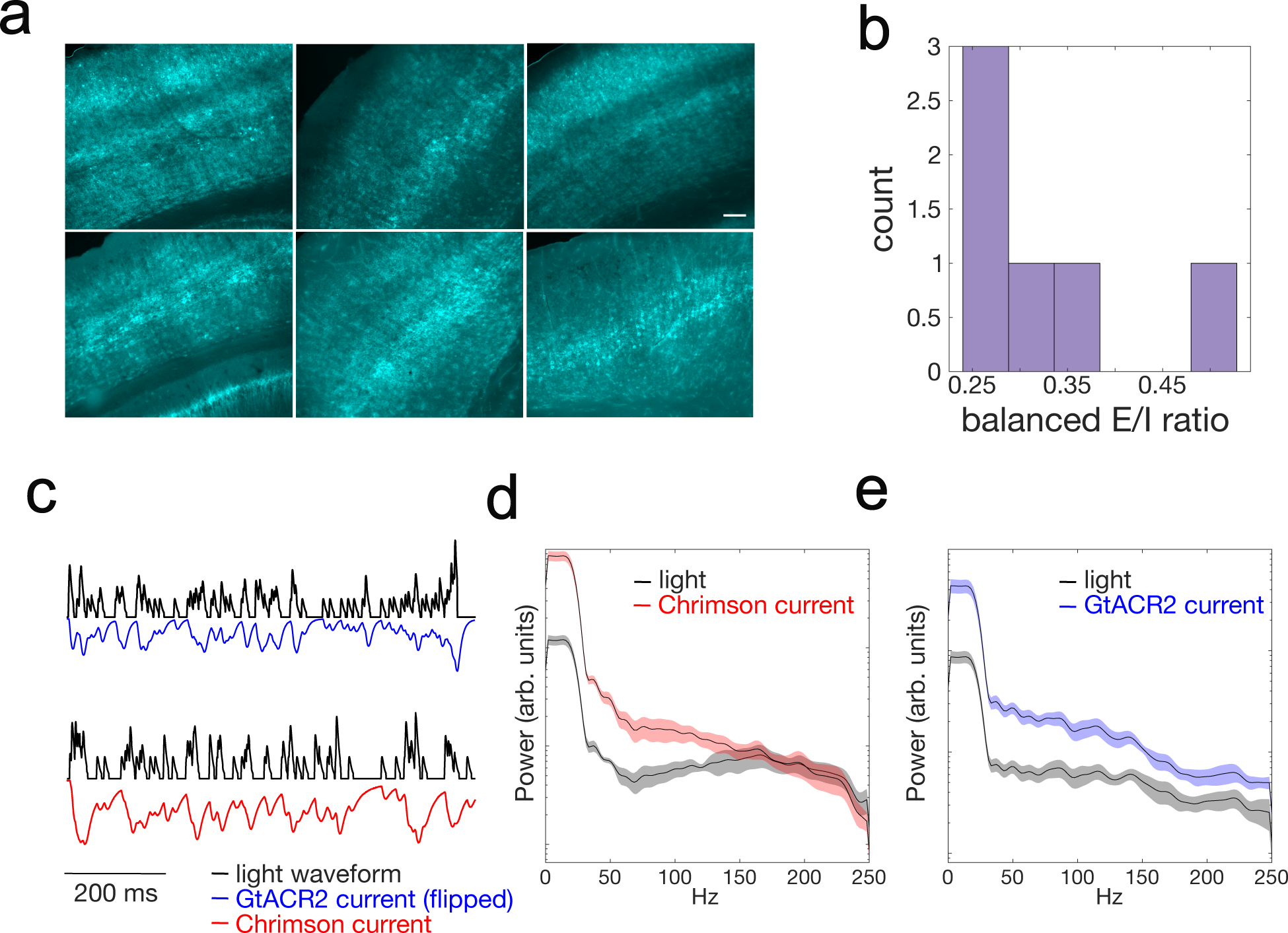
**a)** Example images of somBiPOLES-mCerulean expression in mouse primary visual cortex (V1) in six different mice. Scale bar: 100μm. **b)** Histogram of the operationally balanced blue:red light ratio for BiPOLES-expressing cells in brain slices. **c)** Example command waveforms sent to the blue (top) and red (bottom) LED drivers (black) and the resulting voltage-clamped optogenetically induced conductances. Blue: GtACR2, measured at 0 mV; Red: Chrimson, measured at -70 mV. **d)** Average power spectral density of the light command waveform (black) and the optogenetically induced Chrimson conductance (red). n = 6 cells. **e)** As in d), but for GtACR2 in the same cells.

**Extended Data Figure 2:**
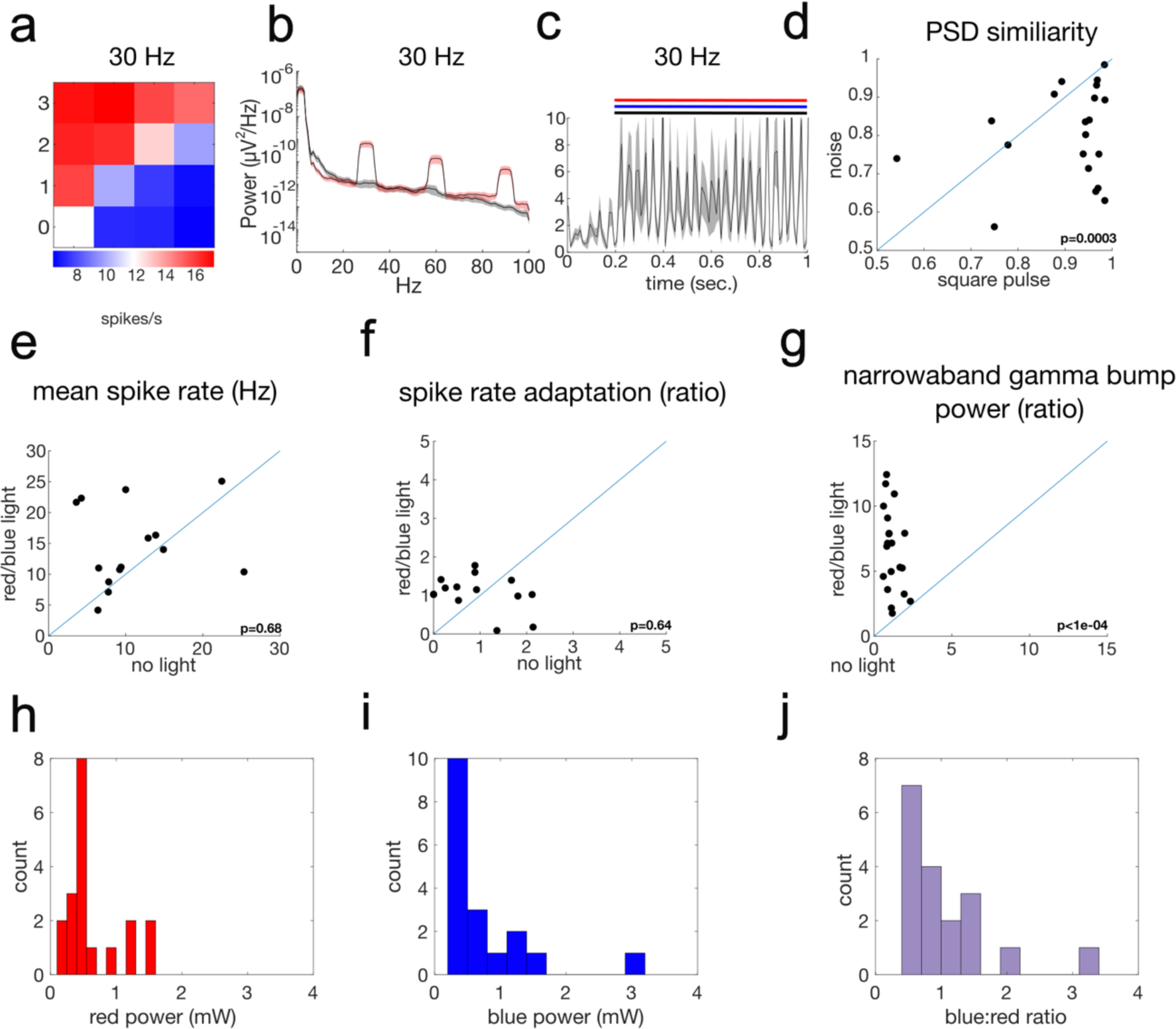
**a)** Average firing rates of juxtacellularly recorded units across light power levels for illumination with 30 Hz sinusoids. n= 20. **c)** Power spectral density plots between control (no light, gray) and maximum red and blue light power for the 30 Hz sinusoidal illumination. Error bars are s.e.m. **c)** PSTH for the recorded units during 30 Hz illumination. **d)** Scatter plot of the cosine similarity of the average power spectral density function between control (light off) and maximum red and blue. **e-g)** Scatter plot of the mean spike rate, spike rate adaptation ratio, and gamma power ratio comparing the control (no light) condition and during illumination with 30 Hz sinusoids of red and blue light. N=20 recordings. **h)** Histogram of the mean red illumination power used in the *in vivo* experiments. **i)** Histogram of the mean blue illumination power used in the *in vivo* experiments. **j)** Histogram of the blue:red ratio for the data present in d)-g).

**Extended Data Figure 3:**
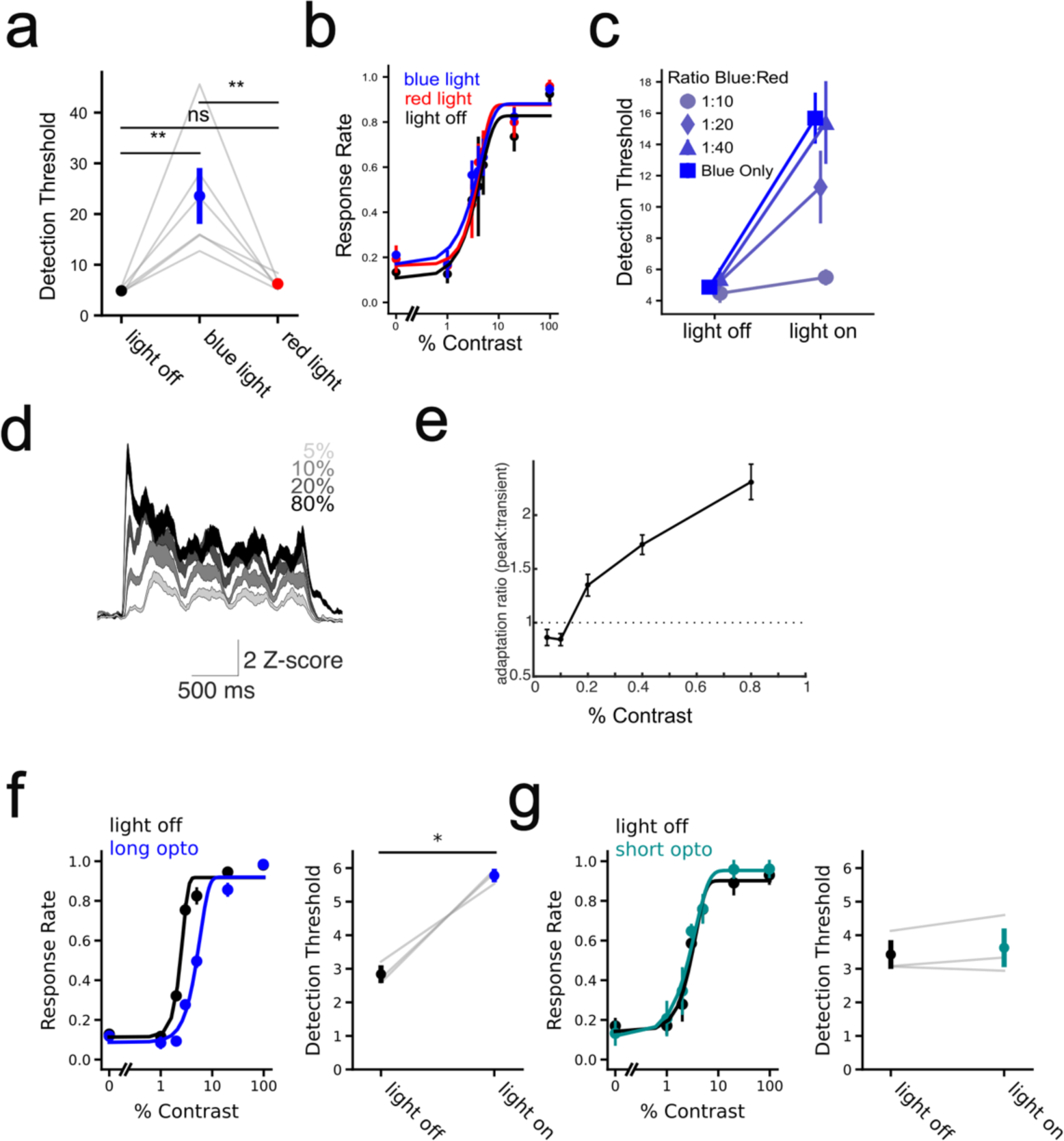
**a)** Detection thresholds in BiPOLES-expressing mice between control (light off, black), blue light only (blue), and red light only (red); p=0.03, paired t-test, n=6 mice, 1 session each. **b)** Average psychometric functions for three mice that did not express BiPOLES. Black: no light, red: red light, blue: blue light. **c)** Detection thresholds in BiPOLES-expressing mice between control (light off) condition and when illuminating with different ratios of blue and red light. n = 6 mice. **d)** Average PSTHs of visually driven spiking across multiple contrasts, demonstrating a strong adapting peak at high contrast, but not at low contrasts; n = 22 mice, 341 units. **e)** Plot of the ratio of mean neural activity at the initial transient response (60-140 ms after visual stimulus onset) to the later more sustained component (220-600 ms after the visual stimulus onset). **f)** Left: Average psychometric functions for mice with no light (black) or blue light during the entire visual stimulus period and response window (blue). Right: Detection thresholds for light off and light conditions. n = 3 mice, p < 0.011, paired t-test. **g)** As in f) except for sessions where the blue light was switched off 100 ms after visual stimulus onset. N = 3 mice; p **=** 0.37, paired t-test.

